# Individual versus fixed parametrization of an electrocutaneous warning signal during manual work tasks

**DOI:** 10.64898/2026.09.17.749350

**Authors:** Eva-Maria Dölker, Rami Farouk, Fatema Altaeh, Jannik Sander, Jens Haueisen

**Affiliations:** Institute of Biomedical Engineering and Informatics, Technische Universität Ilmenau, Ilmenau, Germany

**Author notes:** These authors contributed equally to this work (Rami Farouk, Fatema Altaeh).

**Keywords:** electrocutaneous stimulation, occupational safety, TENS, warning signals, work tasks

## Abstract

Electrocutaneous stimulation provides a means to warn workers of potential hazards. Our previous parameter studies were conducted at rest or under controlled external influences such as vibration, temperature, and humidity, whereas actual work tasks have not yet been addressed. As a step towards greater practical applicability, we conducted two studies in which a circumferential electrocutaneous warning signal was applied to the upper right arm while participants performed a reading task, screw-driving, and polishing. In study 1 (*n* = 32), the pulse interval was adjusted individually to evoke a vibrating sensation, and the intolerance threshold was determined. In study 2 (*n* = 29), a fixed pulse interval of 36 ms was applied to all participants, the warning threshold was determined in addition to the intolerance threshold, and muscle twitches were documented both by participant report and by an independent observer. Study 1 demonstrated that electrocutaneous warning during work tasks is feasible, with a median intolerance threshold higher during polishing (16.5 mA) than during reading or screw-driving (14 mA each). In study 2, the fixed parameter setting elicited the intended pulsating or vibrating sensation at rest in 89 % of the participants in both presentations. The median warning threshold was 8 mA and did not differ between the three work tasks, whereas the intolerance threshold was lower during reading (18 mA) than during screw-driving and polishing (20 mA each). Muscle twitches were largely absent at the warning threshold and became frequent towards the intolerance threshold, with a ventral dominance of their location in both studies. An individual parametrization is therefore not required for the majority of users when a binary warning is to be conveyed, and operating the system slightly above the warning threshold provides a usable amplitude range while minimizing muscle twitches. Future work will focus on electrode optimization and on the objective measurement of motor responses.

## Introduction

In safety-critical work environments, the ability to reliably alert personnel to imminent hazards is paramount — particularly when acoustic [1] or visual [2] cues may be compromised by high noise levels or limited visibility. The audibility of acoustic warning signals is further reduced by hearing protection [3]. In this context, electrocutaneous stimulation emerges as a compelling and innovative modality, offering a promising alternative to conventional warning signal systems [4–10]. In the long term, our objective is to establish a wearable, electrocutaneous warning platform based on textile-integrated electrodes. Such a system should be capable of conveying situationally relevant information by systematically modulating stimulation parameters — including amplitude, frequency, and spatio-temporal patterns — to enable a versatile and intuitive warning modality.

Building on this long-term vision, we defined several overarching requirements. The warning signal must be reliably perceivable, clearly distinguishable, and capable of encoding a small set of categories. Electrode placement should remain comfortable and unobtrusive, avoiding sweat accumulation, motion restriction, interference with other protective gear, or unintended muscle activation interfering with the working tasks. The system must adhere to the EN 60601-1 safety standard [11] and be user-friendly — well tolerated, lightweight, wearable for up to 8 hours, seamlessly integrated into work garments, and supported by robust electronics and power management.

Noninvasive electrical stimulation is used in a wide range of applications, from muscle stimulation [12–17] and transcutaneous electrical nerve stimulation (TENS) [18–20] to more specialized electrocutaneous techniques like acupoint stimulation [21, 22], neuromodulation in neurodegenerative disorders [23] or investigations of locomotor biomechanics [24]. Research on muscle stimulation highlights its relevance for understanding torque generation [13], sensory integration [12], and muscle metabolism [14], as well as for the treatment of neuromotor impairments [16]. A systematic review further emphasizes its benefits for maintaining muscle and bone health in paralyzed limbs [15]. TENS has been widely examined for pain relief [19, 25] and even for reducing postoperative wound infections [20]. Electrocutaneous stimulation is also used in prosthetics to provide sensory feedback [26–28]. However, the design and functionality of electrocutaneous stimulation differ notably in warning systems, as these require higher-amplitude signals [5] than those used for prosthetic sensory feedback. Limit values for electrocutaneous stimulation in warning systems do not exist, so that systematic parameter studies are necessary to develop a reliable electrical warning signal.

A previous parameter study (*n* = 81) [5] investigated the influence of technical stimulation parameters on electrosensory thresholds. Electrodes with an area of 25 mm *×* 40 mm were recommended, along with lateral placement on the upper right arm to avoid muscle twitching, and a pulse width of 150 µs for biphasic rectangular stimulation pulses. Within the same study [7], pulse intervals, amplitudes, and electrode positions were systematically varied. This resulted in pulse interval ranges for the temporal perception categories “single pulses”, “pulsating”, “vibrating”, and “continuous” as well as an almost linear relationship between presented and perceived amplitude and a high discriminability of electrode positions as a basis for the design of electrotactile warning signals. In a pilot study [6] with *n* = 16 participants, a circumferential, vibrating stimulation pattern on the upper right arm was found to induce the highest levels of alertness and was therefore used as warning signal for subsequent studies. Across previous studies [4–7, 9, 10], muscle twitches could not be completely avoided. Therefore, one study (*n* = 15) [8] examined the influence of current flow direction (vertical, diagonal, horizontal). Horizontal stimulation slightly reduced muscle twitch incidence for single pulses, while twitch thresholds showed no systematic differences. For circumferential warning signals, neither muscle twitch occurrence nor attention differed between current directions, indicating equal suitability for electrotactile warning patterns.

In further investigations, electrosensory thresholds were determined as a function of mechanical vibration (*n* = 94) as well as temperature and humidity (*n* = 52) [10]. Increasing vibration amplitude and frequency led to higher thresholds, while female participants exhibited lower values and fewer muscle twitches. Changes in temperature showed only minimal effects. Variations in humidity had no effect on electrosensory thresholds.

Dölker et al. [4] compared textile cuff electrodes with conventional TENS electrodes (*n* = 30) and found overall comparable sensory characteristics, with slightly elevated attention and intolerance thresholds, indicating general suitability for wearable warning systems despite remaining skin–electrode interface impedance issues. In two follow-up studies (*n* = 66) [9], improved textile electrodes exhibited lower perception thresholds and fewer muscle twitches than TENS electrodes, while confirming their suitability for electrotactile warning applications; however, occasional increases in electrode-skin interface impedance were still observed.

Previous studies investigated electrotactile stimulation at rest or under controlled vibration, while the application of electrotactile warning signals during realistic work tasks has not yet been addressed. To close this gap, two separate experimental studies were conducted to evaluate the perception of an electrocutaneous warning signal and the occurrence of muscle twitches during representative work tasks. The first study (*n* = 32) followed an individualized approach, in which the pulse interval of the warning signal was adjusted individually for each participant, whereas the second study (*n* = 29) applied a fixed pulse interval that was identical for all participants and additionally assessed the warning threshold. In both studies, participants performed typical simple work tasks, including reading, operating a cordless screwdriver, and using a polishing machine.

This manuscript is structured as follows. The Methods section describes the participant groups of both studies, the electrocutaneous stimulation setup, the applied work tasks, and the experimental protocols, including the individual and the fixed parametrization of the warning signal, followed by the statistical analysis. The Results section reports the thresholds and the occurrence of muscle twitches for both studies. These findings are interpreted in the Discussion section. Finally, conclusions are drawn and implications for the design of electrotactile warning systems as well as directions for future work are outlined.

## Methods

### Study groups

The descriptive statistics of the two study groups are listed in Table 1, where *n* = 4 participants attended both studies. Participants were recruited between 28 May 2025 and 4 August 2025 for study 1 and between 10 October 2025 and 26 November 2025 for study 2. The ethics committee of the Faculty of Medicine of the Friedrich Schiller University Jena, Germany, approved the studies (reference number 4487-07/15). All methods were carried out in accordance with relevant guidelines and regulations. All participants gave written informed consent.

**Table 1.** Descriptive statistics of the study groups. Age and arm circumference are given as mean *±* standard deviation.

| Property | Study 1: Electrical warning during work tasks with individually adjusted pulse intervals | Study 2: Electrical warning during work tasks with a fixed pulse interval |
| --- | --- | --- |
| Participants | $n = 32$ | $n = 29$ |
| Gender | female: 4, male: 28 | female: 18, male: 11 |
| Age | $29 \pm 7$ years | $28 \pm 8$ years |
| Handedness | right-handed: 30, left-handed: 2 | right-handed: 29, left-handed: 0 |
| Arm circumference (upper right arm) | $32 \pm 5$ cm | $31 \pm 5$ cm |

On the day before and the day of the experiment, participants were instructed to obtain sufficient sleep, abstain from caffeine, nicotine, and alcohol, maintain adequate hydration (approximately 2 l), avoid strenuous physical activity or sports, and refrain from applying skin cream to the upper arms.

## Experimental setup

### Electrical stimulation

The experimental setup was identical to that in our previous publications [4–10]. A brief overview is provided here, while further details can be found in [5]. An in-house program implemented in LabVIEW 2017 (National Instruments, Austin, TX, USA) was used to control a constant-current stimulator DS5 (Digitimer Ltd, Letchworth Garden City, UK) and a multiplexer D188 (Digitimer Ltd, Letchworth Garden City, UK), which activated one of eight output channels to deliver the stimulation signal to the selected electrode pair.

### Electrode configuration

Fourteen reusable self-adhesive TENS electrodes (25 mm *×* 40 mm; axion GmbH, Leonberg, Germany) were arranged in seven pairs along the longitudinal centerline between the shoulder joint and the elbow of the right arm. Electrode pairs were evenly spaced circumferentially at intervals of one eighth of the arm circumference, with one electrode placed 5 mm above and one 5 mm below the centerline. Pairs were numbered consecutively, with pair 1 located ventrally, pair 3 laterally, pair 5 dorsally, and pair 7 medially, and arranged vertically as an upper and a lower electrode. Electrode pair no. 8 (medial–ventral) was omitted due to frequent muscle twitches at this site [5, 6, 9] and because it has been shown to be not relevant for warning pattern presentation [6].

### Warning signal

Based on a previous pilot study [6], a circumferential electrical stimulation pattern around the arm (electrode pairs 1 to 7) that elicits either a pulsating or vibrating sensation, depending on the selected pulse interval, was found to induce the highest levels of alertness and was therefore chosen as the warning signal. The circumferential warning signal was applied sequentially from electrode pair 1 to 7, with each pair stimulated for 0.4 s and separated by a 0.1 s pause. The stimulation signal consisted of biphasic rectangular current pulses with a pulse width of 150 µs. The number of pulses per electrode pair depended on the pulse interval. In contrast to previous studies [6, 8–10], the stimulation amplitude was not derived from single-pulse thresholds, as this approach led to inter-individual inconsistencies in the alerting effect. Instead, the amplitude was increased stepwise after each complete circumferential cycle until sufficient alertness was induced or the stimulation became intolerable. The step size and the maximum amplitude differed between the two studies and are given in the respective paradigm sections. Limit values for electrocutaneous stimulation in warning systems do not exist, and the maximum amplitude was therefore chosen conservatively in both studies, so that the charge density remains more than an order of magnitude below the established limits for safe charge and charge density [29, 30].

### Work tasks

To present the warning signal during work-related activities, three tasks were implemented: a reading task, a cordless screwdriver task, and a polishing machine task.

#### Reading task

Participants were instructed to sit comfortably on a chair and read a short story (see Fig 1a and S1 PDF). To avoid an additional cognitive load from reading in a foreign language, the text was provided in the participant’s native language. The story, entitled “The Disappeared Professor”, was therefore available in Arabic, Simplified Chinese, Croatian, English, Farsi, French, German, Greek, Hindi, Portuguese, Russian, Spanish, and Turkish. The story was generated and translated using ChatGPT (OpenAI, San Francisco, CA, USA) in May 2025. It was designed to be engaging but not overly complex, to have a reading duration of approximately five minutes, and to be easily translatable across languages. After reading had commenced, the presentation of the warning signal was initiated with a short temporal delay of about 5 s.

**Fig 1.**
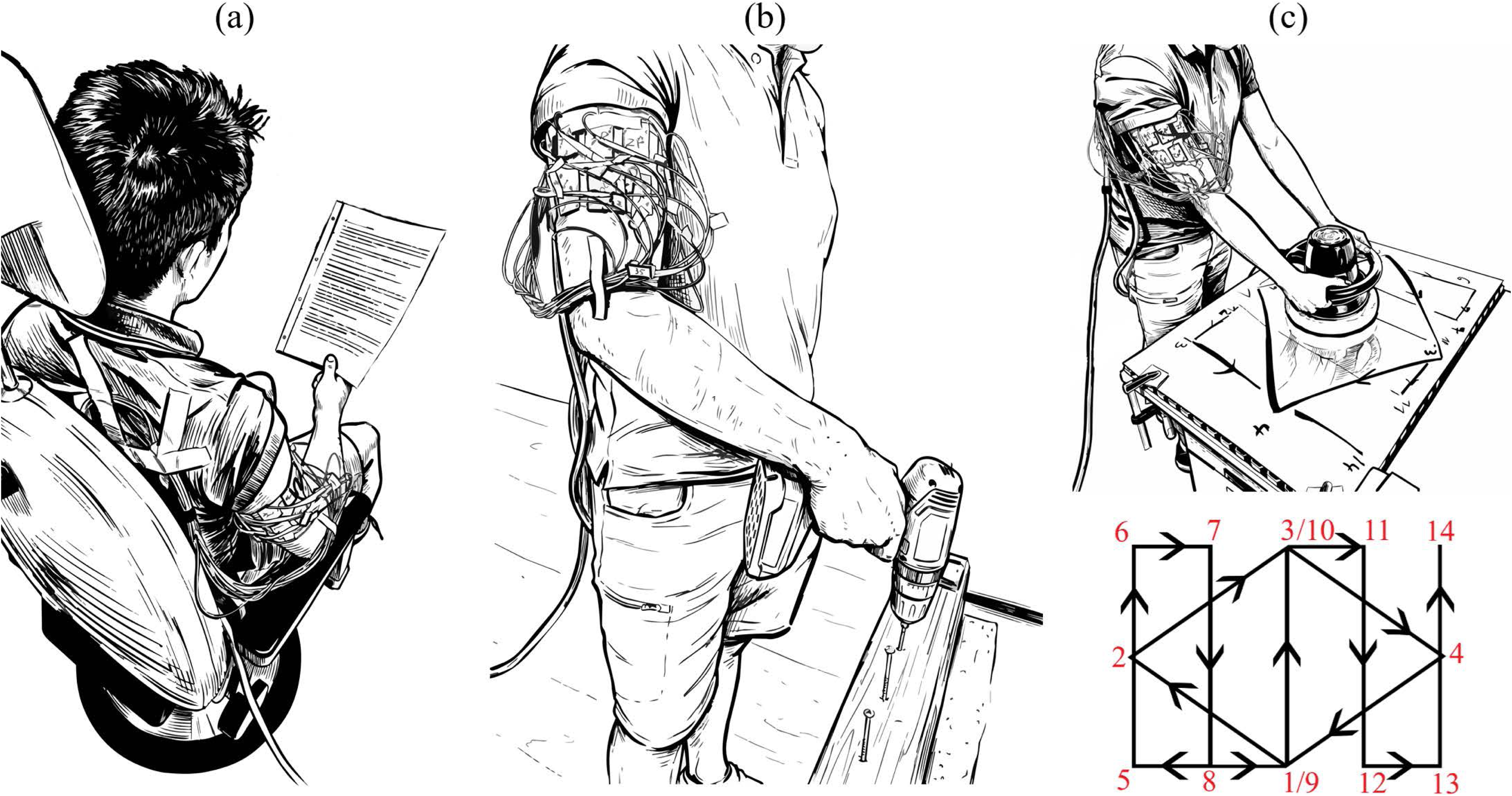
Work tasks. (a): Reading task. (b): cordless screwdriver task. (c): polishing machine task (top) and sketch of the prescribed movement pattern for the polishing task (bottom). For this preprint, the original photographs were replaced by drawings.

#### Cordless screwdriver task

Participants were required to drive eight screws into a wooden beam with their right arm, using a cordless screwdriver BOSCH EasyImpact 18V-40 (Robert Bosch GmbH, Gerlingen-Schillerhöhe, Germany). The task is shown in Fig 1b. After all screws had been inserted, they were removed and subsequently reinserted. The presentation of the warning signal was initiated with a short temporal delay of about 5 s.

#### Polishing machine task

Participants were required to guide a polishing machine (Cartrend GmbH, Rülzheim, Germany) at 3000 revolutions per minute over an aluminum plate with both arms, following a predefined movement pattern (see Fig 1c). The polishing machine was placed at point 1 and then moved across the aluminum plate in the sequence 1–2–3–4–1–5–6–7–8–9–10–11–12–13–14, after which it was returned to point 1 and then the pattern was repeated. The presentation of the warning signal was initiated with a short temporal delay of about 5 s.

### Preparation

Both studies followed the same preparation procedure. The devices were switched on 30 min before the experiment to ensure adequate warm-up. In addition, the DS5 current stimulator was stabilized by connecting a 1 kΩ load resistor and repeatedly applying stimulation pulses. To optimize the electrode-skin interface impedance, the participant’s right upper arm was cleaned with ethanol and subsequently moistened using axion electrode contact spray (250 ml, axion GmbH, Leonberg, Germany). The experiment was conducted at a room temperature of approximately 23 °C.

### Experimental paradigm of study 1

#### Reference threshold determination

To allow the participant to become accustomed to the electrical stimulation, three single pulse thresholds were determined at the lateral electrode pair 3. These thresholds were additionally used to determine the individual pulse interval that elicited a vibrating sensation. Based on previous studies, the defined thresholds were: (1) the perception threshold *A*_p_ at which the stimulus is just perceptible for the first time; (2) the muscle twitch threshold *A*_m_ corresponding to the onset of visible muscle twitches; and (3) the intolerance threshold *A*_i_, which elicits a stimulus perceived as intolerable. To determine these threshold values, a single biphasic rectangular stimulation pulse with a pulse width of 150 µs was applied. The current amplitude was increased stepwise from 0 mA up to a maximum of 25 mA, using increments of 0.1 mA, 0.2 mA, or 0.5 mA. The steps were chosen adaptively by the trained operator. The operator asked the participant about the spatial perception at all three thresholds and about the qualitative perception at the perception and the intolerance threshold. For the qualitative perception, there were the following choices in the questionnaire: knocking, scratching, stinging, pain, muscle twitch, tickling, itching, pinching, squeezing. The choices for the spatial perception included: in the area of the stimulated electrode pair, at another electrode pair or at another part of the body.

The threshold determination procedure was repeated ten times, which allowed the participant to identify each threshold more reliably. The reported single pulse thresholds are the mean of the last three repetitions.

#### Individual pulse interval determination

In order to determine the individual pulse interval at which a vibration sensation was perceived, ten biphasic rectangular stimulation pulses were presented to electrode pair 3 with a pulse interval of 19 ms and the previously determined mean perception threshold *A*_p_. The current was increased, if necessary, in steps of 1 mA until the stimulus could be clearly perceived. If the perceived sensation was not described as vibrating, the pulse interval was shortened when the sensation was described as pulsating and lengthened when it was described as continuous, with the step size chosen by the trained operator [7]. The individual pulse interval was documented and subsequently served as the basis for the applied warning signal.

#### Electrical warning during work tasks

The warning signal was presented during the three work tasks described above, always performed in the same order: reading, operating a cordless screwdriver, and operating a polishing machine. After the respective task had commenced, the presentation of the warning signal was initiated with a short temporal delay of about 5 s. Starting from 1 mA, the stimulation amplitude was increased in steps of 1 mA after each complete circumferential cycle up to a maximum of 25 mA or until the stimulation was perceived as intolerable, at which point it was stopped and the corresponding intolerance threshold *A*_i_ was noted. In addition, any occurring muscle twitching was quantified with respect to its intensity (“perceptible”, “visible”, “arm movement”) and its location (electrode pair number, upper arm, other location).

### Experimental paradigm of study 2

#### Warning signal without individual parametrization

In our previous studies [8–10] and in study 1, the pulse interval at which a participant perceives a vibrating sensation was determined individually. The main aim of study 2 was to realize and evaluate an electrocutaneous warning signal without individual parametrization. Building on study 1, the paradigm was extended in two respects. The pulse interval was fixed across participants, and the warning threshold was assessed in addition to the intolerance threshold.

To determine a suitable fixed pulse interval, a previous study [7] with *n* = 81 participants was re-evaluated, of whom 80 completed the experiment on varying pulse intervals. In that study, five consecutive biphasic rectangular pulses with a pulse width of 150 µs and pulse intervals from 200 ms down to 0.5 ms were presented to each participant, and the pulse interval ranges for the temporal sensations “single pulses”, “pulsating”, “vibrating” and “continuous” were determined. Fig 2 shows these ranges for each participant.

**Fig 2.**
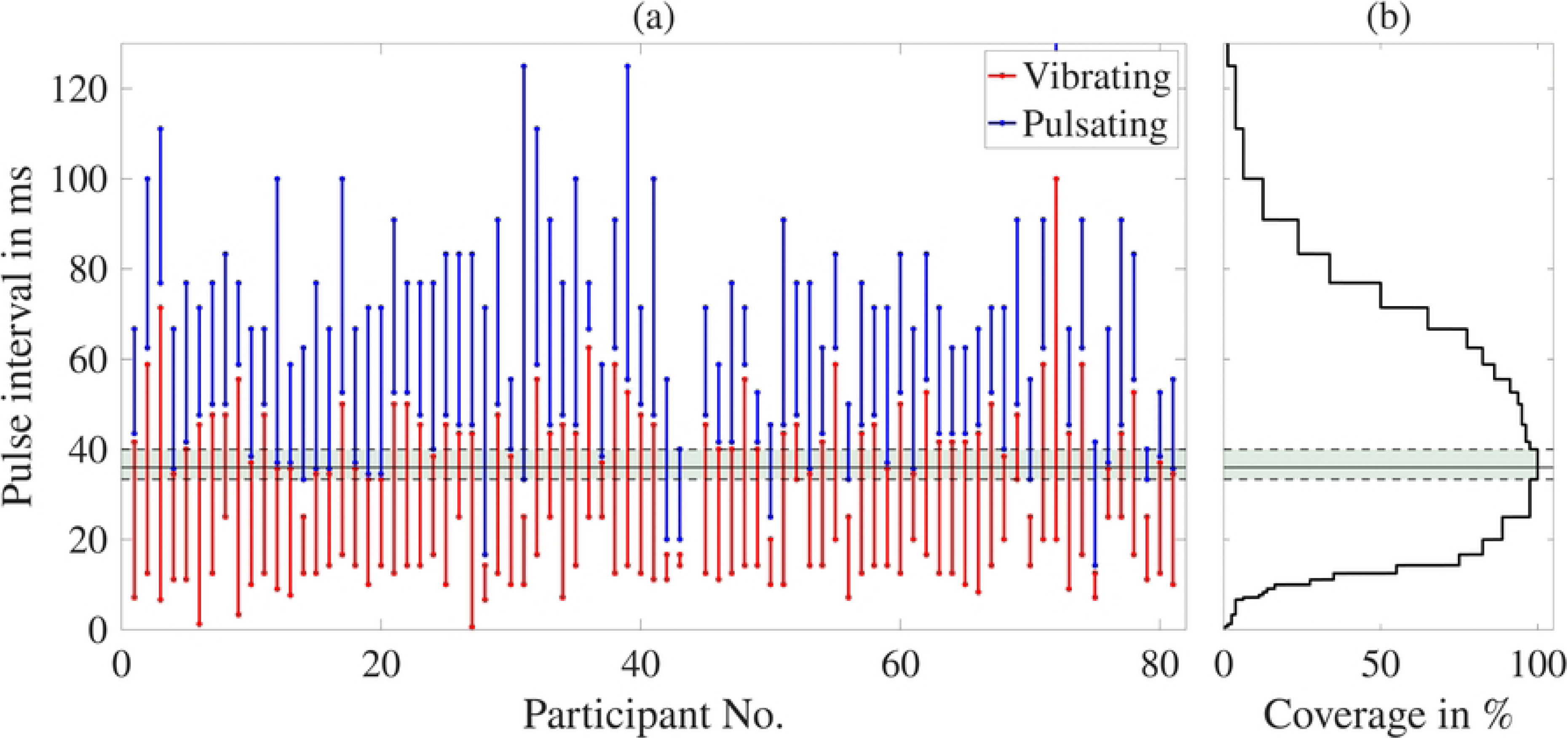
Individual pulse intervals of. *n* = 80 **participants from a previous study [7].** (a): Individual pulse interval ranges. Red ranges indicate the pulse intervals perceived as a vibrating sensation, blue ranges those perceived as a pulsating sensation. For participant no. 72, the maximum pulse interval reached 200 ms. The axis is truncated to a range of 0 to 130 ms for better visibility. For participant no. 44, the experiment on varying pulse intervals could not be carried out, so that 80 of the 81 participants of the previous study are shown. (b): Proportion of participants whose combined pulsating and vibrating range covers the respective pulse interval. The dashed lines mark the lower (33 ms) and upper (40 ms) boundary of the range covered by all 80 participants, and the shaded area between them the plateau of 100 % coverage. The solid black line indicates the pulse interval of 36 ms applied to all participants in study 2. Apparent gaps between the red and the blue range of a participant span exactly one step of the tested pulse intervals in [7] and therefore reflect the resolution of the stimulation protocol rather than a pulse interval range without a temporal sensation.

No single pulse interval was perceived as vibrating by all participants (Fig 2a). It is known from a previous study [6], however, that a circumferential stimulation signal generating a pulsating sensation leads to levels of alertness comparable to a vibrating one. Both temporal qualities are therefore suitable for a warning signal, and we combined them into a single range per participant, reaching from the shortest pulse interval perceived as vibrating to the longest one perceived as pulsating.

In Fig 2a, the vibrating and the pulsating range of a participant never touch. As the pulse interval was varied in discrete steps in the previous study [7], the transition between the two temporal qualities is resolved to one step, which separates the two ranges. We therefore treated the combined range of each participant as continuous.

For each pulse interval, we determined the proportion of participants whose combined range covers it (Fig 2b). The proportion reaches 100 % between 33 ms and 40 ms and decreases outside this range, to 97.5 % at 25 ms and to 96.2 % at 45 ms. Within these boundaries, every participant perceived the stimulation as either pulsating or vibrating. The stability of the boundaries was assessed by omitting each participant in turn and by drawing 10 000 random samples of 80 participants with replacement.

The two boundaries are each defined by two participants, the lower one by participants 52 and 69, the upper one by participants 43 and 79 (see Fig 2a), so that omitting any single participant leaves the range unchanged. The resampling reproduced the observed boundaries in 87 % of the samples. In the remaining 13 %, the lower boundary fell to 25 ms, and the upper boundary rose to at most 45.5 ms. A pulse interval of 36 ms was contained in every sample and was therefore applied to all participants in study 2.

### Electrical warning during rest

The participant was seated in a relaxed position with the right arm resting loosely on the armrest. The aim was to familiarize the participant with the electrocutaneous stimulation and to determine the individual warning and intolerance thresholds under resting conditions. Starting from 2 mA, the stimulation amplitude was increased in steps of 2 mA up to a maximum of 24 mA. The warning threshold was recorded as the stimulation amplitude at which the participant described the stimulation as sufficiently attention-catching to be perceived as a warning, such that an ongoing task would be interrupted in a real scenario. As soon as the stimulation was no longer tolerable, it was stopped immediately and the corresponding amplitude was documented as the intolerance threshold. If the stimulation remained tolerable, the presentation was stopped at 24 mA. In addition, the temporal perception was recorded using the categories “single pulses”, “pulsating”, “vibrating”, and “continuous”. Any occurring muscle twitches were quantified with respect to intensity (“none”, “visible”, “arm movement”) and location (“medial”, “ventral”, “lateral”, “dorsal”, “whole arm”, “other location”). In contrast to study 1, the category “perceptible” was deliberately omitted, as it could not be verified by an external observer. Beyond the participant self-report, the muscle twitches were documented by an independent observer. The observer documented them during the presentation of the warning signal and was instructed not to comment on them, whereas the participant was asked only after the end of the presentation, so that the two assessments were obtained independently of each other. This procedure was performed twice, followed by a break of 5 min.

### Electrical warning during work tasks

The warning signal was presented during the three work tasks, always performed in the same order: reading, operating a cordless screwdriver, and operating a polishing machine. After the respective task had commenced, the warning signal was initiated with a short temporal delay of about 5 s. Starting from 2 mA, the stimulation amplitude was increased in steps of 2 mA up to a maximum of 24 mA. The warning threshold and the intolerance threshold were determined as in the resting condition. If the stimulation remained tolerable, the presentation was stopped at 24 mA. Any occurring muscle twitches were documented with respect to intensity and location by participant self-report and by the independent observer, as in the resting condition.

### Statistics

All statistical analyses and data visualizations were carried out in MATLAB R2025b (The MathWorks, Natick, MA, USA). The thresholds were determined on a coarse amplitude grid and contain a large number of tied values, and the sample sizes were moderate (*n* = 32 in study 1 and *n* = 29 in study 2). Non-parametric methods were therefore used throughout [31]. Comparisons between conditions were carried out as paired Wilcoxon signed-rank tests on the within-participant differences, using the normal approximation with correction for tied ranks. The exact test was not used, as the stepwise variation of the amplitude produces a large number of tied ranks and zero differences, which violates its assumptions. Zero differences were discarded before ranking. Because of their large number, all comparisons were repeated with the procedure proposed by Pratt [32], which retains the zeros during ranking, and the conclusions were unchanged. To account for multiple comparisons, a Bonferroni correction was applied [33]. Each *p*-value was multiplied by the number of tests and evaluated against a significance level of *α* = 0.05. Corrected *p*-values exceeding 1 were truncated to 1.

Threshold and amplitude distributions are reported as the median together with the interquartile range [25th–75th percentile] and illustrated as boxplots, which show the distributions across participants and may therefore overlap even when the within-participant differences are systematic. For the pairwise comparisons of the work tasks, the distributions are additionally shown as half violins, each combined with a boxplot, and the two values of the same participant are connected by a line, so that the direction of the within-participant differences becomes visible. For study 1 and study 2, the strength of muscle twitches is shown as bar charts, whereas the location was quantified as the mean relative proportion per location.

## Results

### Study 1

#### Reference thresholds

Fig 3 shows the distribution of the perception, muscle twitch and intolerance single pulse thresholds of *n* = 32 participants of study 1. For *n* = 14 participants no muscle twitches occurred, and *n* = 27 participants did not reach the intolerance threshold within 25 mA. The median thresholds [IQR] are 4.03 mA [3.45 mA–4.55 mA] for the perception threshold, 20 mA [20 mA–25 mA] for the muscle twitch threshold and 25 mA [25 mA–25 mA] for the intolerance threshold.

**Fig 3.**
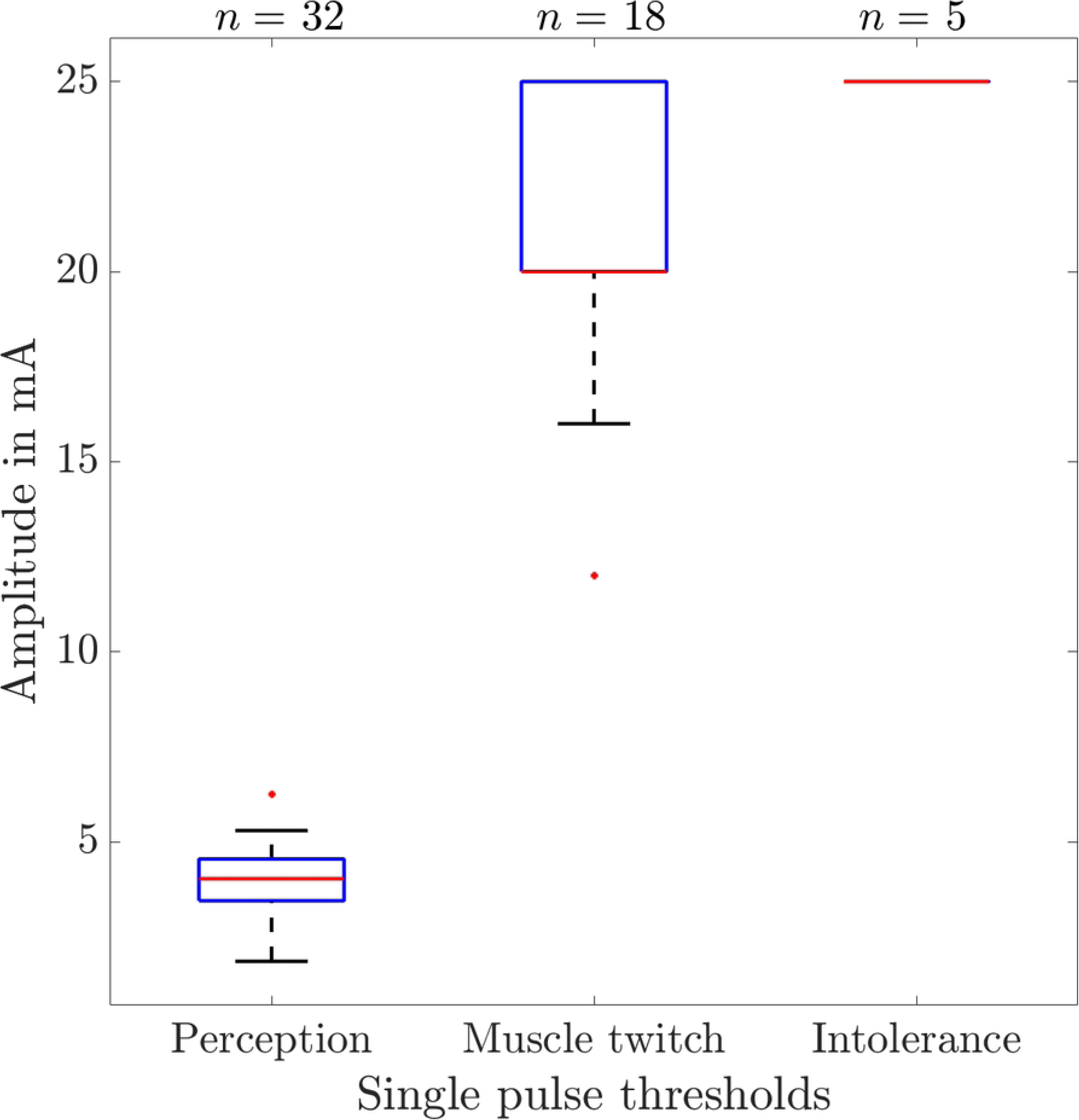
Single pulse thresholds of study 1. Perception, muscle twitch and intolerance single pulse thresholds of *n* = 32 participants. For *n* = 14 participants no muscle twitches occurred, and *n* = 27 participants did not reach the intolerance threshold within 25 mA.

The paired Wilcoxon signed-rank tests on the within-participant differences revealed a significant difference between the perception and the muscle twitch threshold (*p ≪* 0.05). The remaining pairwise comparisons showed no significant differences, which reflects the small number of participants who reached the intolerance threshold (*n* = 5 and *n* = 3 for the pairwise complete cases) rather than an absence of a difference. The *p*-values are Bonferroni corrected. All 32 participants reported a knocking sensation at the perception threshold. Of the *n* = 5 participants who reached the intolerance threshold, 4 reported a stinging and 1 a knocking sensation. The spatial sensation was localized to the area of the stimulated electrode pair for all participants. Muscle twitches occurred in 18 of 32 participants. In 72 % of these cases, the twitching occurred at electrode pair 3, in 16 % at electrode pair 4, in 6 % between electrode pairs 3 and 4, and in 6 % at electrode pair 5. No muscle twitches were observed outside of the area of the electrodes.

#### Electrical warning during work tasks

The median of the individually determined pulse intervals for a vibrating sensation was 19 ms [17.5 ms–19 ms]. Fig 4 shows the pairwise comparisons of the intolerance thresholds during the presentation of the electrocutaneous warning signal for the three work tasks reading, operating a cordless screwdriver and operating a polishing machine. For *n* = 4, *n* = 7 and *n* = 8 out of 32 participants the intolerance threshold was not reached. The median values [IQR] of the intolerance thresholds were 14 mA [10.5 mA–18 mA] during reading, 14 mA [12 mA–20 mA] during screw-driving and 16.5 mA [14 mA–19 mA] during polishing.

**Fig 4.**
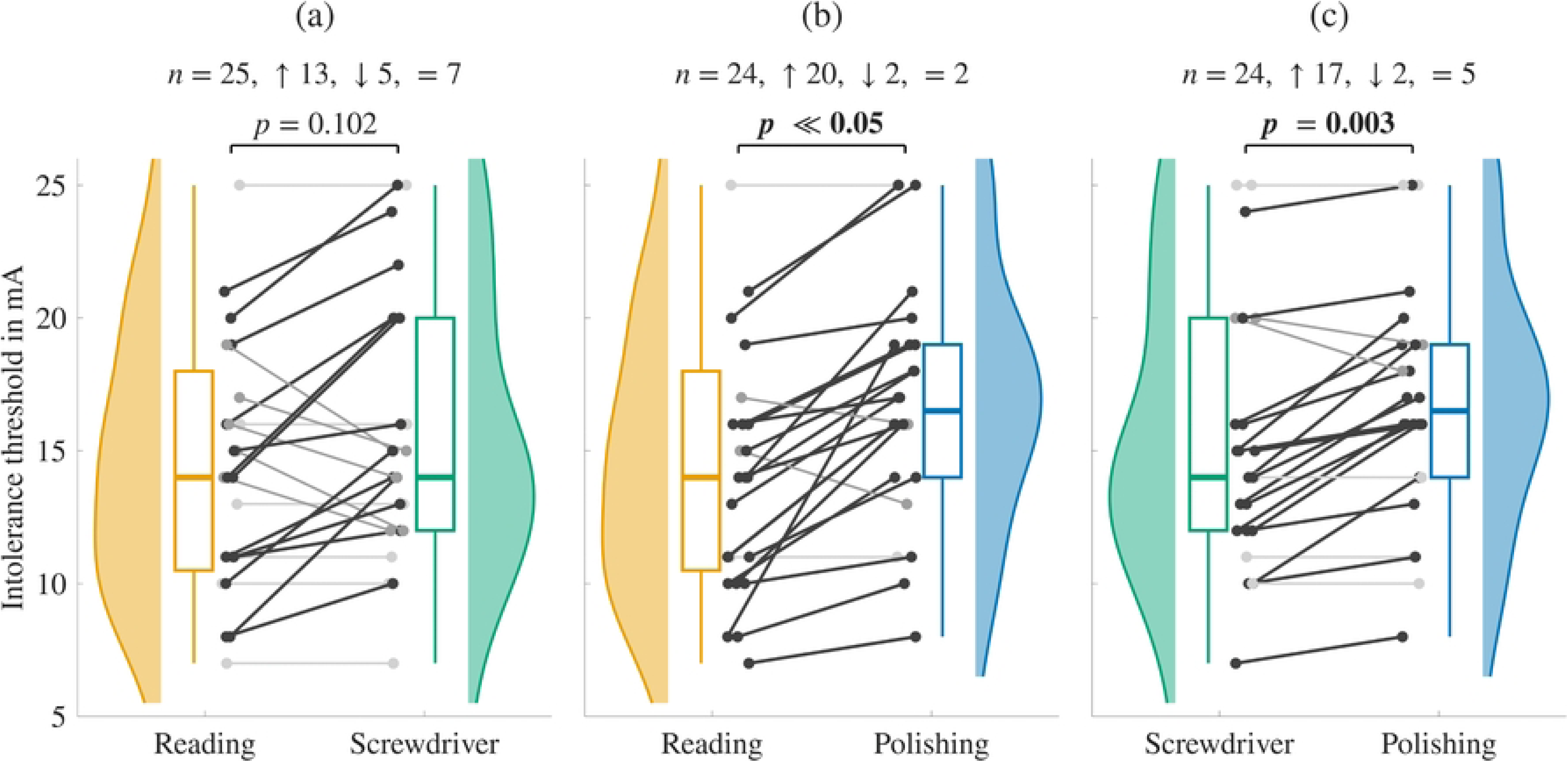
Intolerance thresholds of the applied warning signal of study 1. During the work tasks reading, operating a cordless screwdriver, and operating a polishing machine *n* = 4, *n* = 7 and *n* = 8 out of 32 participants did not reach the intolerance threshold. The half violins and boxplots show all participants who reached the intolerance threshold in the respective task (*n* = 28, *n* = 25 and *n* = 24 for reading, screw-driving and polishing). The connecting lines and the statistical tests are based on the pairwise complete cases, whose number is given above each panel.

Participants who did not reach the intolerance threshold within 25 mA were excluded pairwise, leaving 25, 24 and 24 of the 32 participants for the comparisons reading vs. cordless screwdriver, reading vs. polishing machine and cordless screwdriver vs. polishing machine. These numbers are close to the numbers per task, because the censoring is nested: every participant who did not reach the intolerance threshold during reading did not reach it during screw-driving and polishing either. The paired Wilcoxon signed-rank tests on the within-participant differences revealed no significant difference between reading and operating a cordless screwdriver (*p* = 0.102), but significant differences between reading and operating the polishing machine (*p ≪* 0.05) and between operating the cordless screwdriver and operating the polishing machine (*p* = 0.003). The *p*-values are Bonferroni corrected. The within-participant difference was zero for 7, 2 and 5 participants, so that 18, 22 and 19 participants remained. Among these, the intolerance threshold was higher during the second of the two tasks in 13, 20 and 17 cases.

Fig 5 shows the strength of the muscle twitches observed during the presentation of the warning signal.

**Fig 5.**
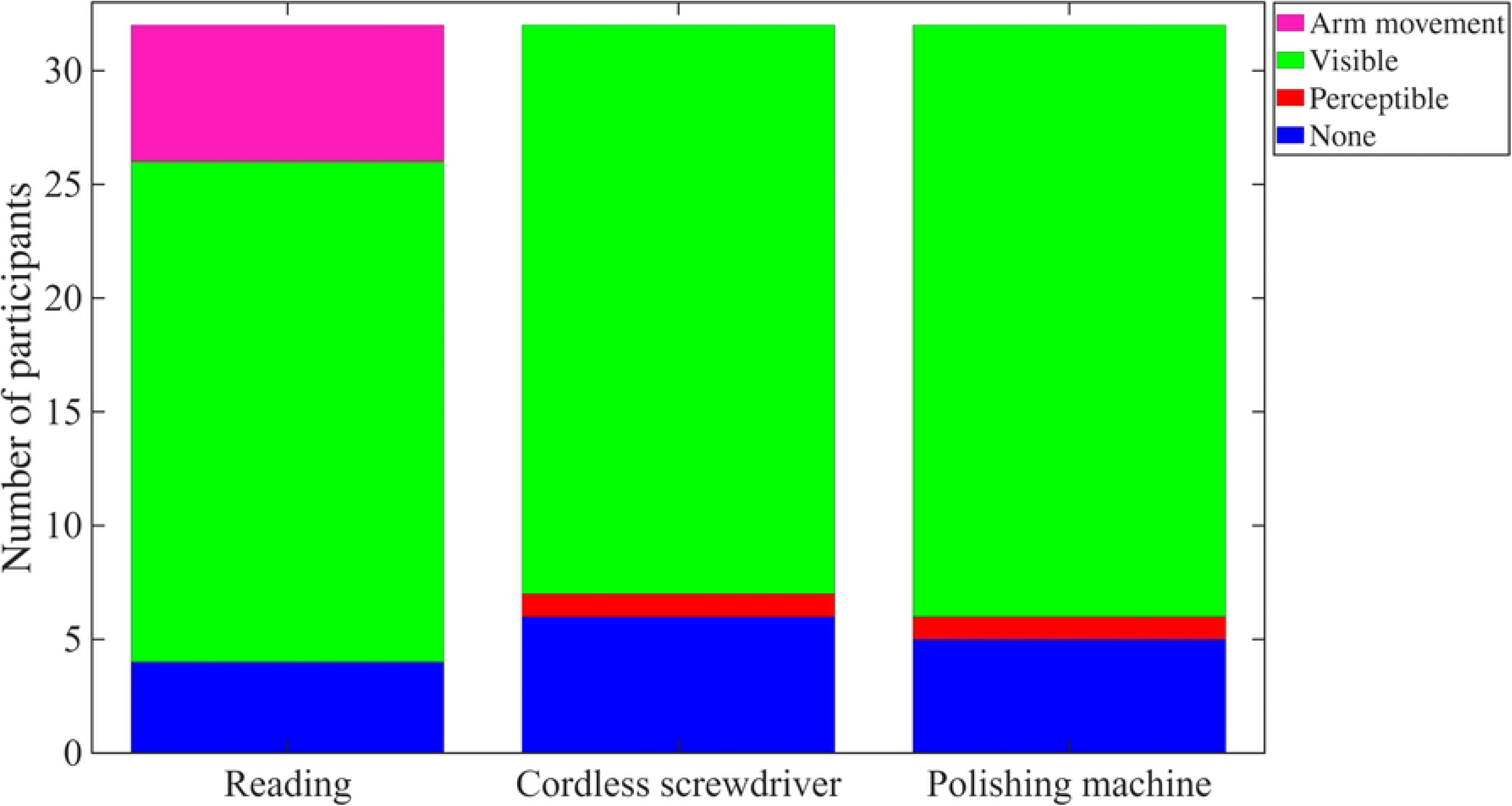
Muscle twitch strengths during the applied warning signal of study 1. The warning signal was presented during the work tasks reading, operating a cordless screwdriver, and operating a polishing machine. The muscle twitches were observed by the researcher conducting the experiment, with the category “perceptible” based on the participant report, and refer to the whole amplitude range up to the intolerance threshold.

Muscle twitches occurred in 88 %, 81 % and 84 % of the participants during reading, operating the cordless screwdriver and operating the polishing machine. During reading, 69 % showed visible muscle twitches and 19 % showed arm movements. During the two tool-based tasks, no arm movements occurred. The muscle twitches were visible in 78 % and 81 % of the participants, and one participant in each of these two tasks showed a muscle twitch that was perceptible but not visible.

The muscle twitch locations documented in study 1 as electrode pair numbers were mapped onto the regions “medial”, “ventral”, “lateral”, “dorsal”, “whole arm” and “other location” to ensure comparability with the location categories used in study 2. Following the circumferential arrangement of the electrode pairs, pair 1 was assigned to “ventral”, pair 3 to “lateral”, pair 5 to “dorsal” and pair 7 to “medial”. The pairs 2, 4 and 6 were located between two of these regions. Whenever such an intermediate pair was documented, an adjacent electrode pair had additionally been reported, which resolved the assignment unambiguously in all cases. Twitches reported for the whole upper arm were assigned to “whole arm”, all remaining reports to “other location”.

For the quantification of the muscle twitch location, the relative proportions of the reported locations were calculated for each participant who experienced muscle twitches. If a twitch occurred at only one location, this location was assigned a value of 1. If it occurred at several locations, the value of 1 was distributed evenly over them, so that a report of ventral and dorsal corresponds to the proportions medial: 0, ventral: 0.5, lateral: 0, dorsal: 0.5, whole arm: 0 and other location: 0. The mean of these proportions over all participants with muscle twitches is shown in Fig 6 for the three work tasks.

**Fig 6.**
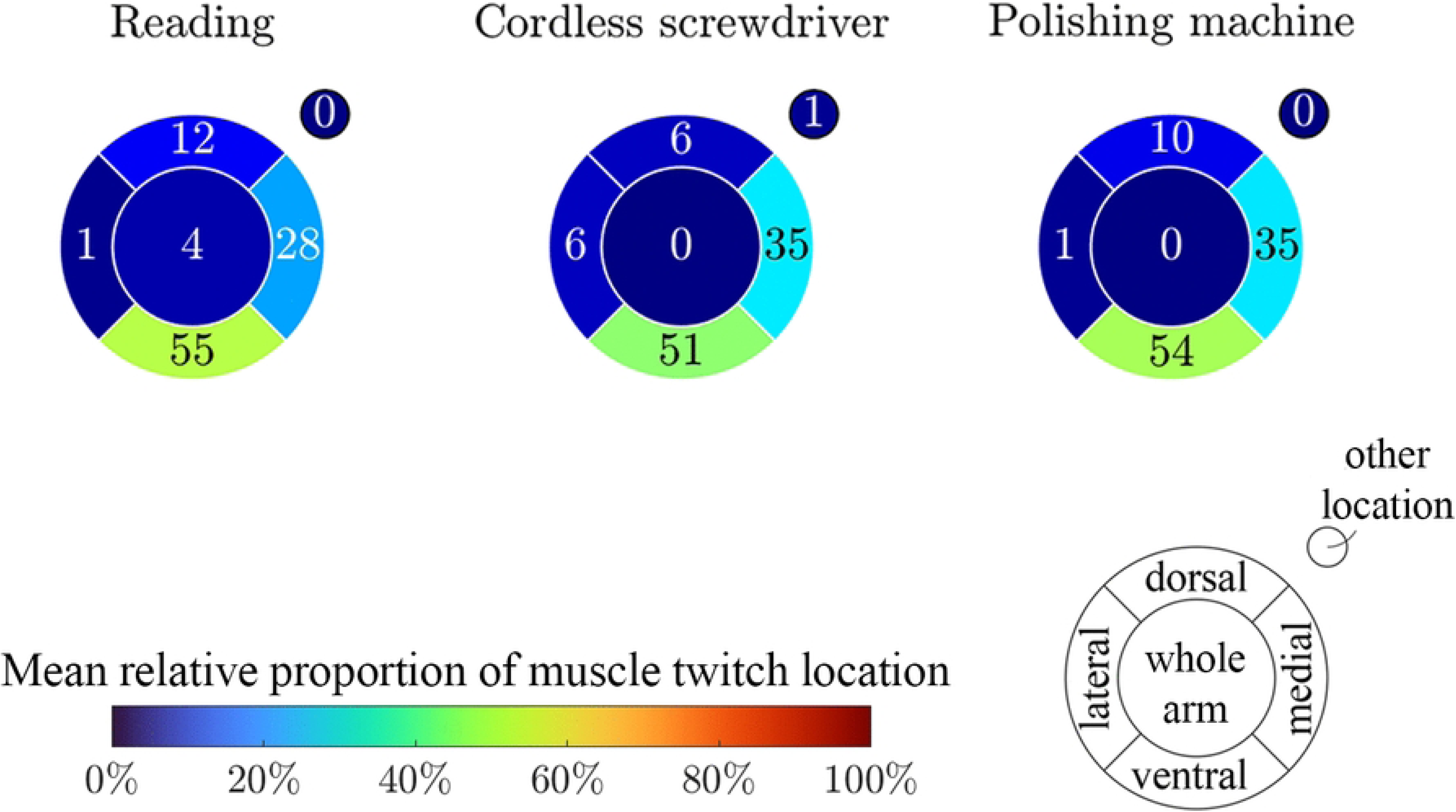
Mean relative proportion of muscle twitch location during the applied warning signal of study 1. The warning signal was presented during the work tasks reading, operating a cordless screwdriver, and operating a polishing machine.

Over 50 % of the muscle twitches occurred at the ventral position of the upper right arm, followed by 28 % to 35 % at the medial position. The muscle twitch locations were comparable between the three tasks. One participant experienced a muscle twitch reaching down to the hand while operating the cordless screwdriver, which was the only report assigned to “other location”.

### Study 2

#### Electrical warning during rest

Fig 7 shows the warning and intolerance thresholds during the first and second presentation of the electrocutaneous warning signal under resting conditions.

**Fig 7.**
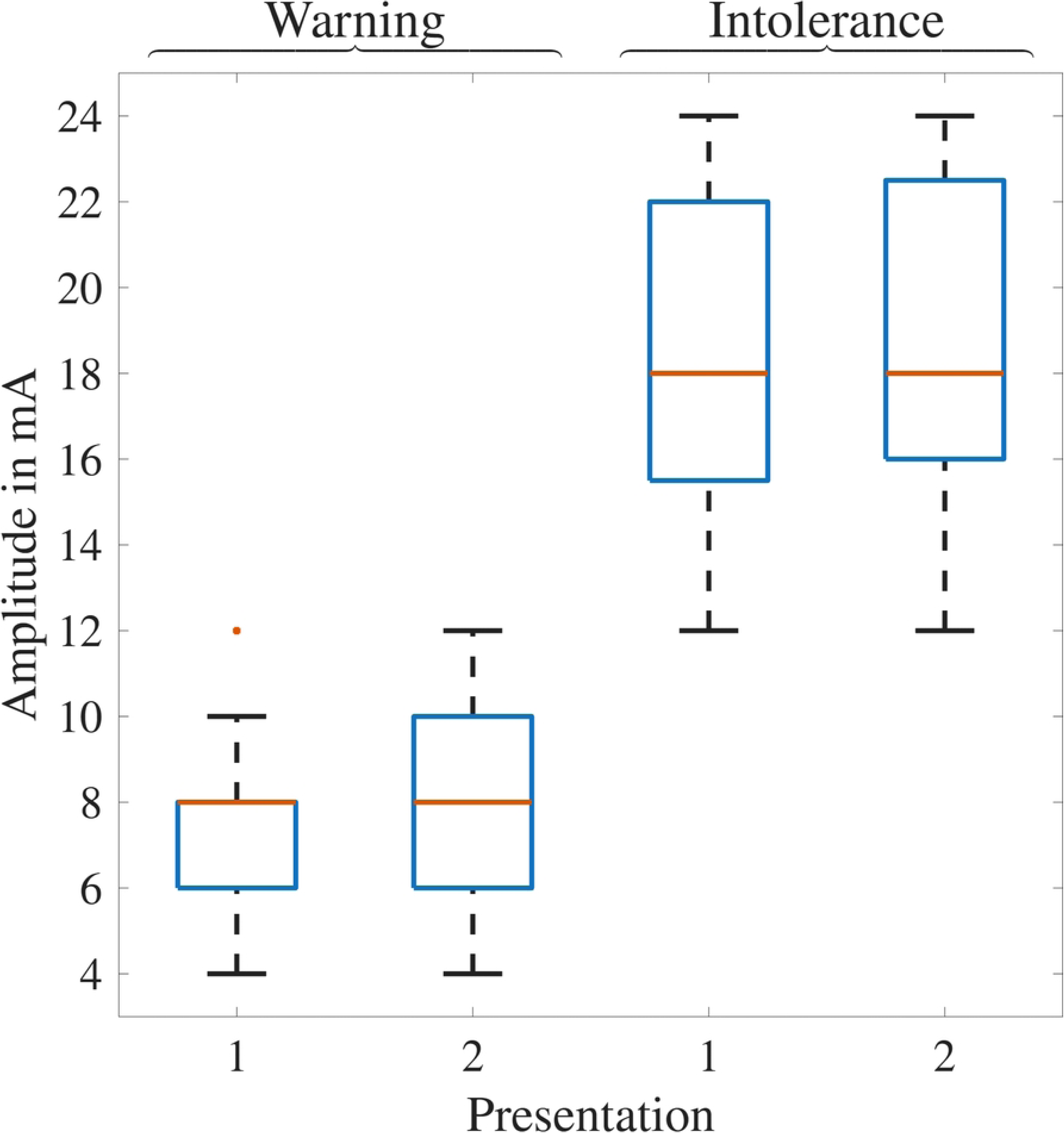
Warning and intolerance thresholds of the applied warning signal during rest of study 2. For *n* = 6 and *n* = 7 out of 29 participants in presentation 1 and 2, the stimulation remained tolerable up to the maximum amplitude of 24 mA, so that their intolerance threshold is a lower bound.

For 21 % and 24 % of the participants the stimulation remained tolerable up to the maximum amplitude of 24 mA during the first and second presentation of the warning signal at rest. The median [IQR] of the warning threshold was 8 mA [6 mA–8 mA] for the first and 8 mA [6 mA–10 mA] for the second presentation. The intolerance threshold was 18 mA [15.5 mA–22 mA] for the first and 18 mA [16 mA–22.5 mA] for the second presentation. The paired Wilcoxon signed-rank tests on the within-participant differences showed no significant difference between the two presentations, neither for the warning threshold (*p* = 1.000) nor for the intolerance threshold (*p* = 1.000). The intolerance thresholds were higher than the warning thresholds (*p ≪* 0.05). The *p*-values are Bonferroni corrected.

Fig 8 shows the temporal perception of the warning signal during the first and second presentation at rest.

**Fig 8.**
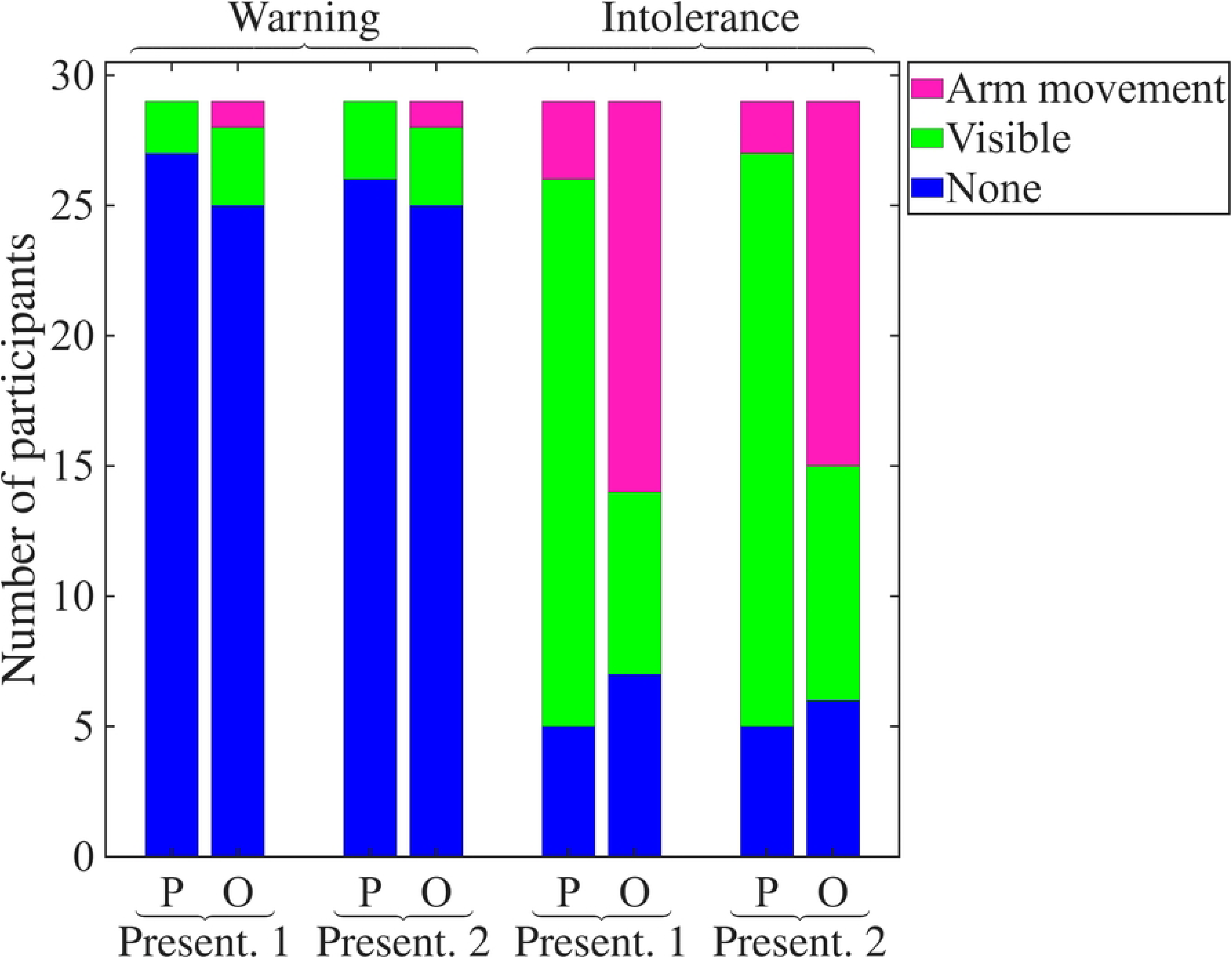
Temporal perception of the applied warning signal of study 2 at rest. The warning signal was presented two times. For one participant no information is available, for another one the information for presentation 2 is missing.

During the first presentation of the warning signal at rest, 3 % of the participants reported a pulsating, 86 % a vibrating and 7 % a continuous perception. For the second presentation, the proportions were 10 %, 79 % and 3 %. For one participant no information on the temporal perception is available, for another one the information for the second presentation is missing.

Fig 9 shows the muscle twitch strengths during the first and second presentation of the warning signal at rest, reported by the participant and documented by the independent observer.

**Fig 9.**
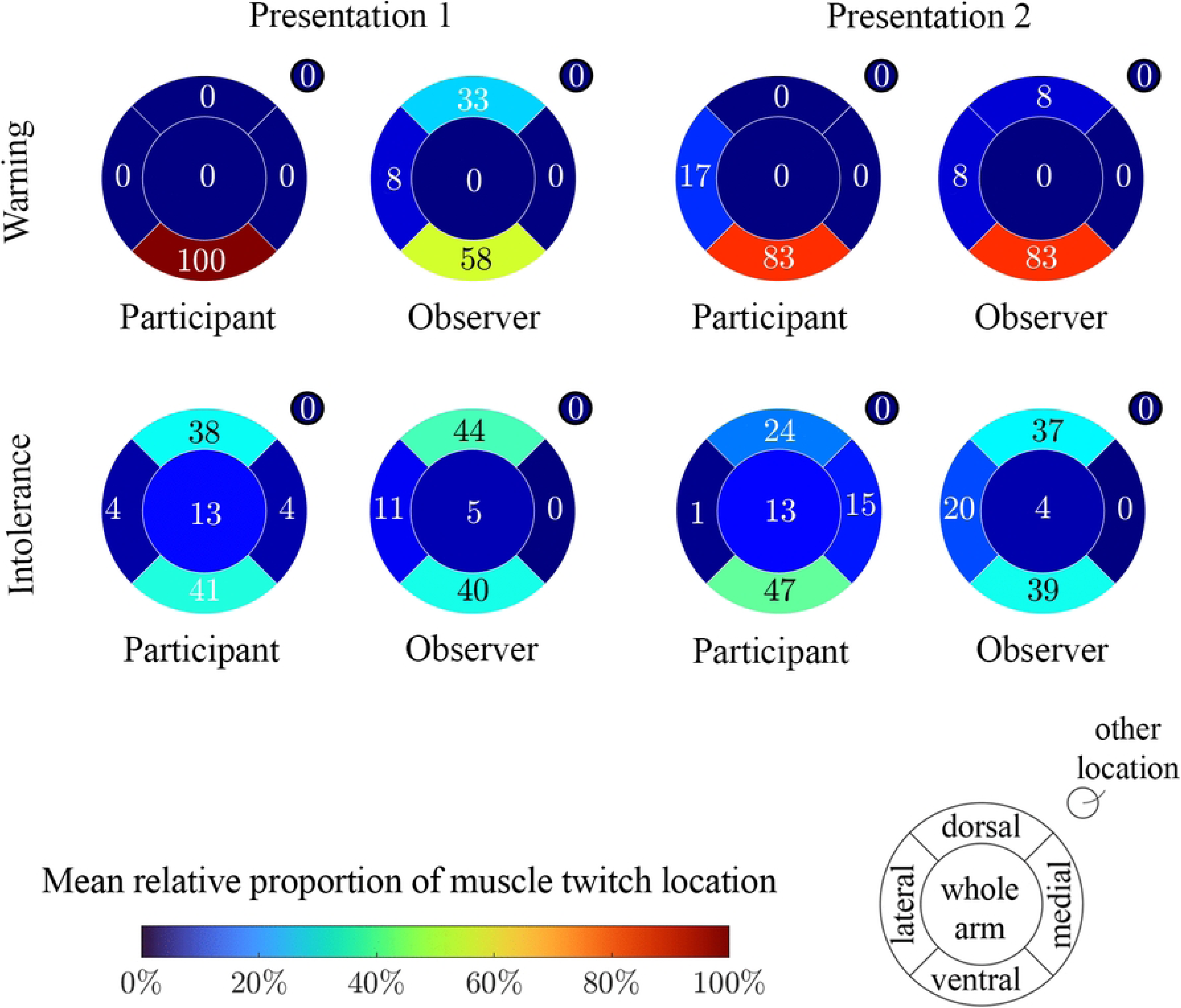
Muscle twitch strengths of study 2 at rest. The strengths are given at the warning and at the intolerance threshold, reported by the participant (P) and documented by the independent observer (O). The warning signal was presented two times (Present. 1 and Present. 2).

The number of participants experiencing muscle twitches increased at the intolerance threshold in comparison to the warning threshold (Fig 9). The distinction between muscle twitching and no muscle twitching was consistent between the participant report and the independent observer. The observer, however, documented a higher proportion of arm movements than the participants reported.

The relative proportions per location were calculated as described for study 1, separately for the warning and the intolerance threshold, for the first and second presentation of the warning signal, and for the participant report and the independent observer (Fig 10).

**Fig 10.**
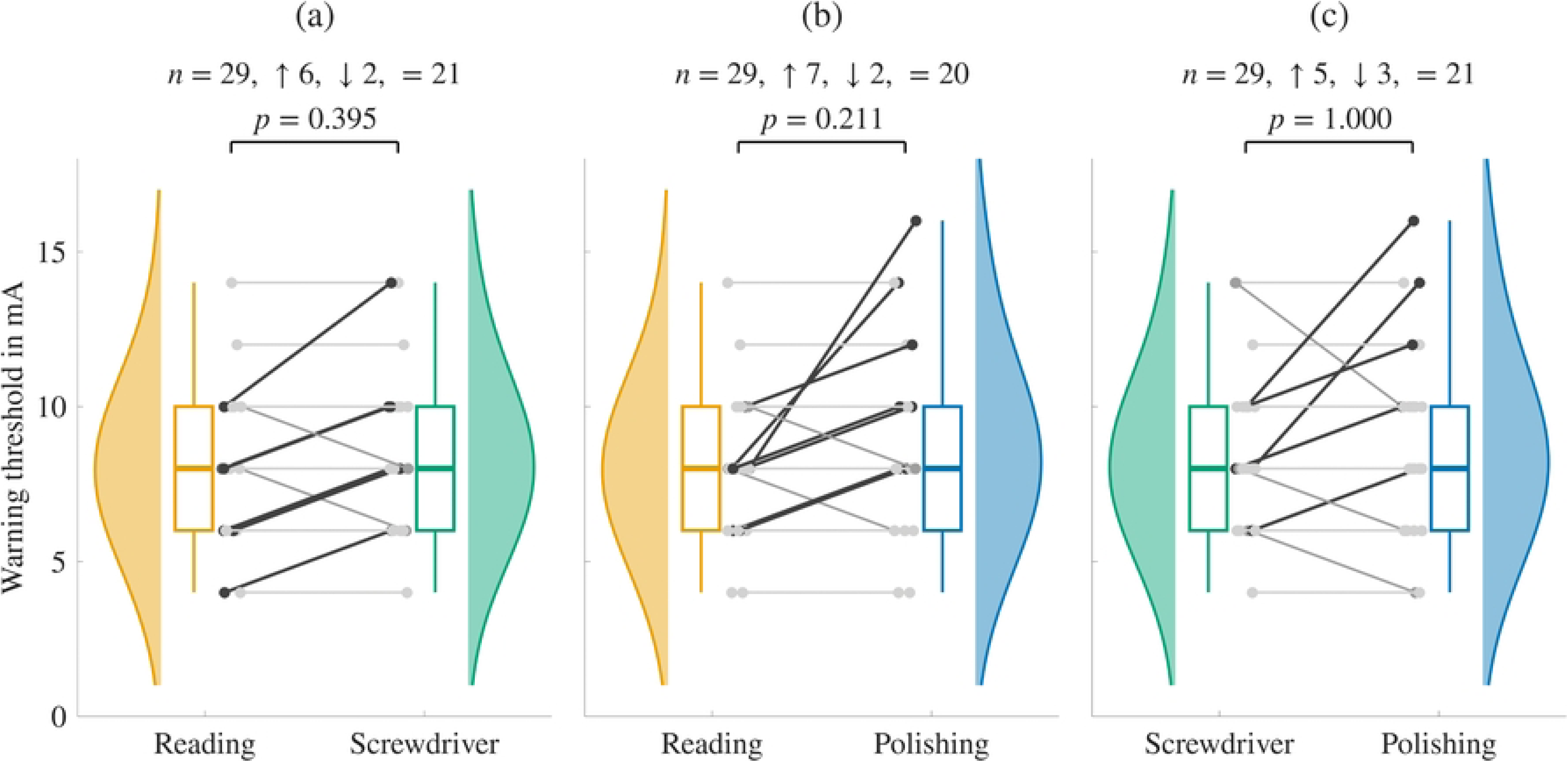
Mean relative proportion of muscle twitch location of study 2 at rest. The proportions are given at the warning and at the intolerance threshold, reported by the participant and documented by the independent observer. The warning signal was presented two times (Presentation 1 and 2). The proportions are averaged over the participants who experienced muscle twitches.

The few participants who already showed muscle twitches at the warning threshold (*n* = 2 and *n* = 3 in the two presentations) reported them mainly in the ventral region of the upper right arm. In accordance with the higher amplitudes of the intolerance thresholds (Fig 7), the proportion of participants with muscle twitches increased (Fig 9) and the location diversified towards dorsal and whole-arm responses (Fig 10). The medial and the lateral region were the least frequent locations.

#### Electrical warning during work tasks

Fig 11 and Fig 12 show the warning and the intolerance thresholds during the three work tasks of study 2.

**Fig 11.**
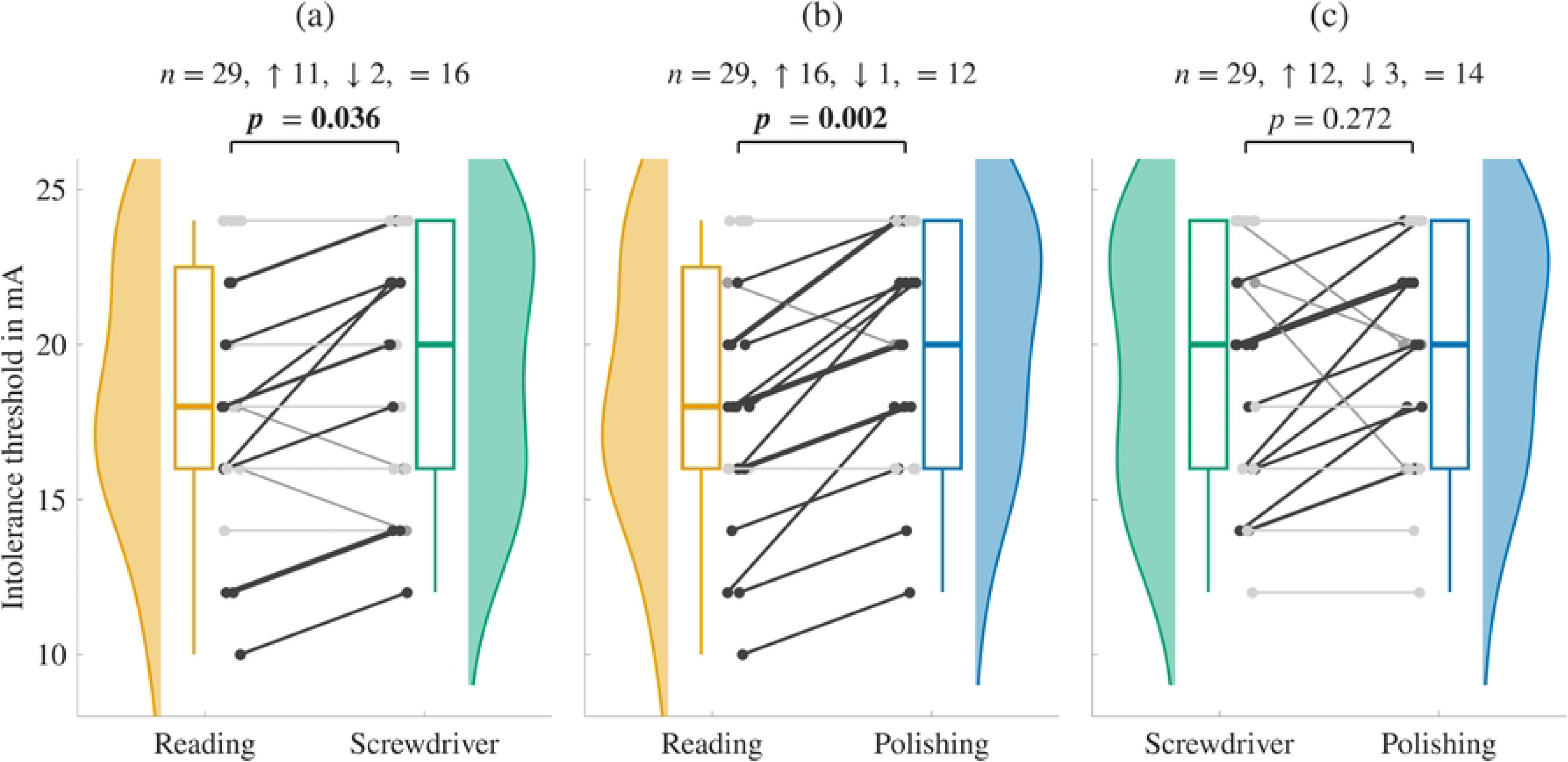
Warning thresholds of the applied warning signal of study 2. Pairwise comparisons of the work tasks reading, operating a cordless screwdriver (Screwdriver) and operating a polishing machine (Polishing). Each task is shown as a half violin representing the smoothed distribution and a boxplot giving median, interquartile range and whiskers. The lines connect the two values of the same participant and are dark where the threshold increases from the left to the right task. Above each panel, *n* is the number of participants, followed by the number of increases, decreases and unchanged values, and the Bonferroni-corrected *p*-value of the paired Wilcoxon signed-rank test.

**Fig 12.**
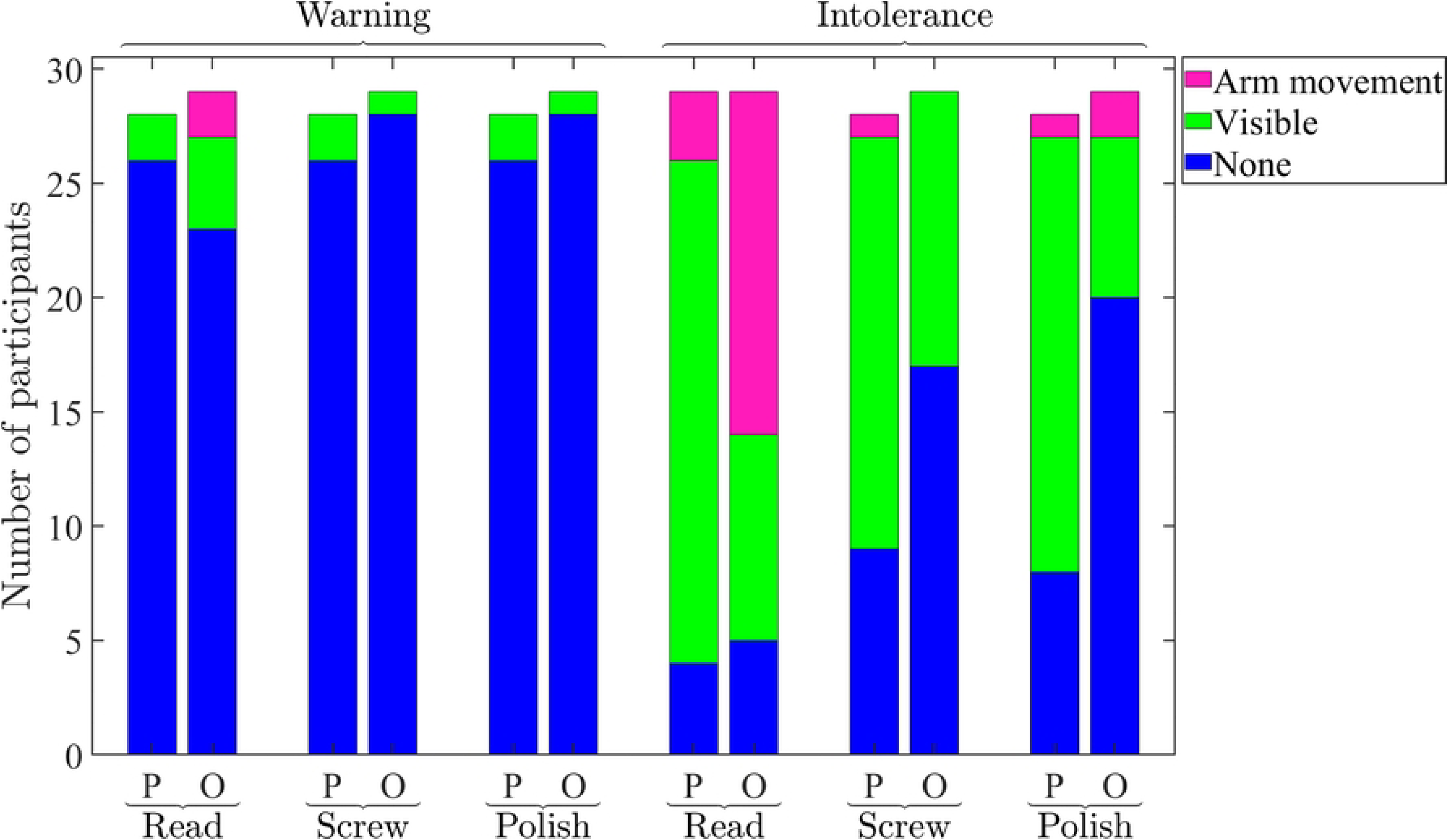
Intolerance thresholds of the applied warning signal of study 2. Pairwise comparisons of the work tasks reading, operating a cordless screwdriver (Screwdriver) and operating a polishing machine (Polishing), presented as in Fig 11. For *n* = 7, *n* = 9 and *n* = 10 out of 29 participants a warning signal presentation up to the maximum value of 24 mA was possible during the three tasks, so that their intolerance threshold is a lower bound and they contribute an unchanged value to every comparison.

The warning thresholds were 8 mA [6 mA–10 mA] during all three work tasks. The paired Wilcoxon signed-rank tests on the within-participant differences showed no significant differences between the work tasks (reading vs. cordless screwdriver: *p* = 0.395, reading vs. polishing machine: *p* = 0.211, cordless screwdriver vs. polishing machine: *p* = 1.000). The intolerance thresholds were 18 mA [16 mA–22.5 mA] during reading and 20 mA [16 mA–24 mA] during screw-driving and polishing. The corresponding tests showed significant differences between reading and screw-driving (*p* = 0.036) and between reading and polishing (*p* = 0.002), whereas the comparison between screw-driving and polishing showed no significant difference (*p* = 0.272). All *p*-values are Bonferroni corrected. For 24 %, 31 % and 34 % of the participants the stimulation remained tolerable up to the maximum amplitude of 24 mA during reading, screw-driving and polishing.

For these participants, 24 mA is a lower bound of the intolerance threshold rather than the threshold itself, and 7 of the 29 participants reached this bound in all three work tasks. They contribute a difference of zero to every comparison, irrespective of whether their intolerance thresholds would have differed between the tasks. Together with the amplitude steps of 2 mA, this results in 16, 12 and 14 of the 29 within-participant differences being zero for the comparisons reading vs. cordless screwdriver, reading vs. polishing machine and cordless screwdriver vs. polishing machine. Among the remaining 13, 17 and 15 participants, the intolerance threshold was higher during the second of the two tasks in 11, 16 and 12 cases.

Fig 13 shows the muscle twitch strengths at the warning and at the intolerance threshold during the three work tasks, reported by the participant and documented by the independent observer.

**Fig 13.**
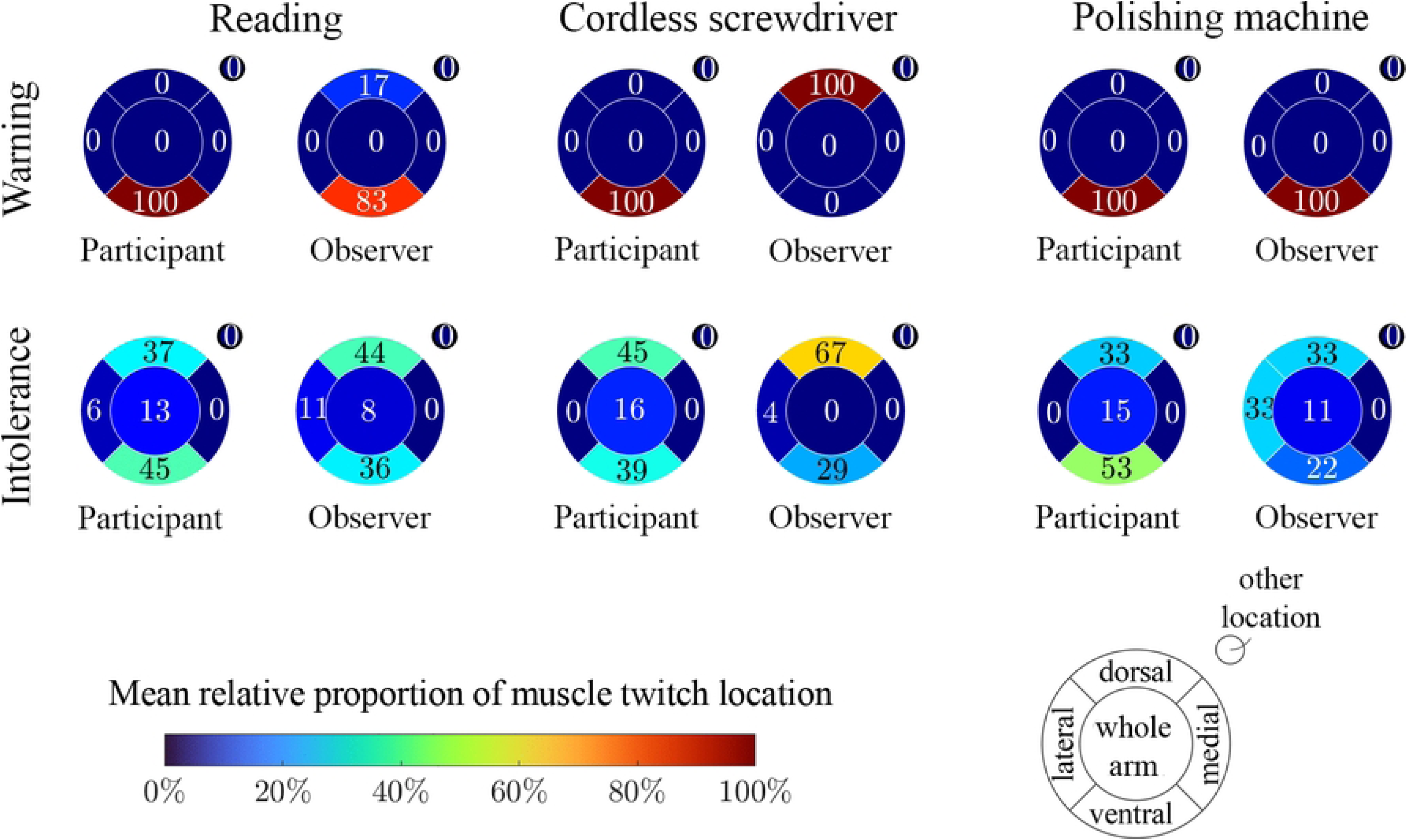
Muscle twitch strengths of study 2 during the work tasks. The strengths are given at the warning and at the intolerance threshold, reported by the participant (P) and documented by the independent observer (O). The warning signal was presented during the work tasks reading (Read), operating a cordless screwdriver (Screw), and operating a polishing machine (Polish).

One statement of the participant on the muscle twitch strength is missing in each of the work task paradigms, except for the intolerance threshold during reading. In the two tool-based tasks, the missing statement belongs to the same participant at both thresholds.

Across all three tasks, 7 % of the participants reported muscle twitches at the warning threshold, compared with 21 %, 3 % and 3 % documented by the observer during reading, screw-driving and polishing (Fig 13). The observer thus confirmed the participant reports in most cases, but documented visible muscle twitches in single participants who had not noticed them themselves. At the intolerance threshold, these proportions increased to 86 %, 68 % and 71 % in the participant reports and to 83 %, 41 % and 31 % in the observer reports. One participant showed a pronation of the lower right arm during reading, and two participants briefly let go of the polishing machine. For reading at the intolerance threshold, the distinction between muscle twitching and no muscle twitching was consistent between the participant report and the observer, but the observer documented a larger proportion of arm movements. During screw-driving and polishing, the proportion of muscle twitches was higher in the participant reports than in the observer reports.

The relative proportions per muscle twitch location were calculated analogously (Fig 14). The few participants who experienced muscle twitches already at the warning threshold reported them mainly in the ventral region of the upper right arm. During screw-driving at the warning threshold, the location was ventral in 100 % of the participant reports and dorsal in 100 % of the observer reports. This is not a disagreement about the location, as the two proportions originate from different participants. Two participants reported ventral muscle twitches that the observer did not see, and the observer documented a dorsal muscle twitch in a third participant who did not notice it himself. With increasing amplitudes at the intolerance threshold, the proportion of participants experiencing muscle twitches increased and the localization diversified towards dorsal, lateral and whole-arm responses. No muscle twitch was localized to the medial region at the intolerance threshold.

**Fig 14.**
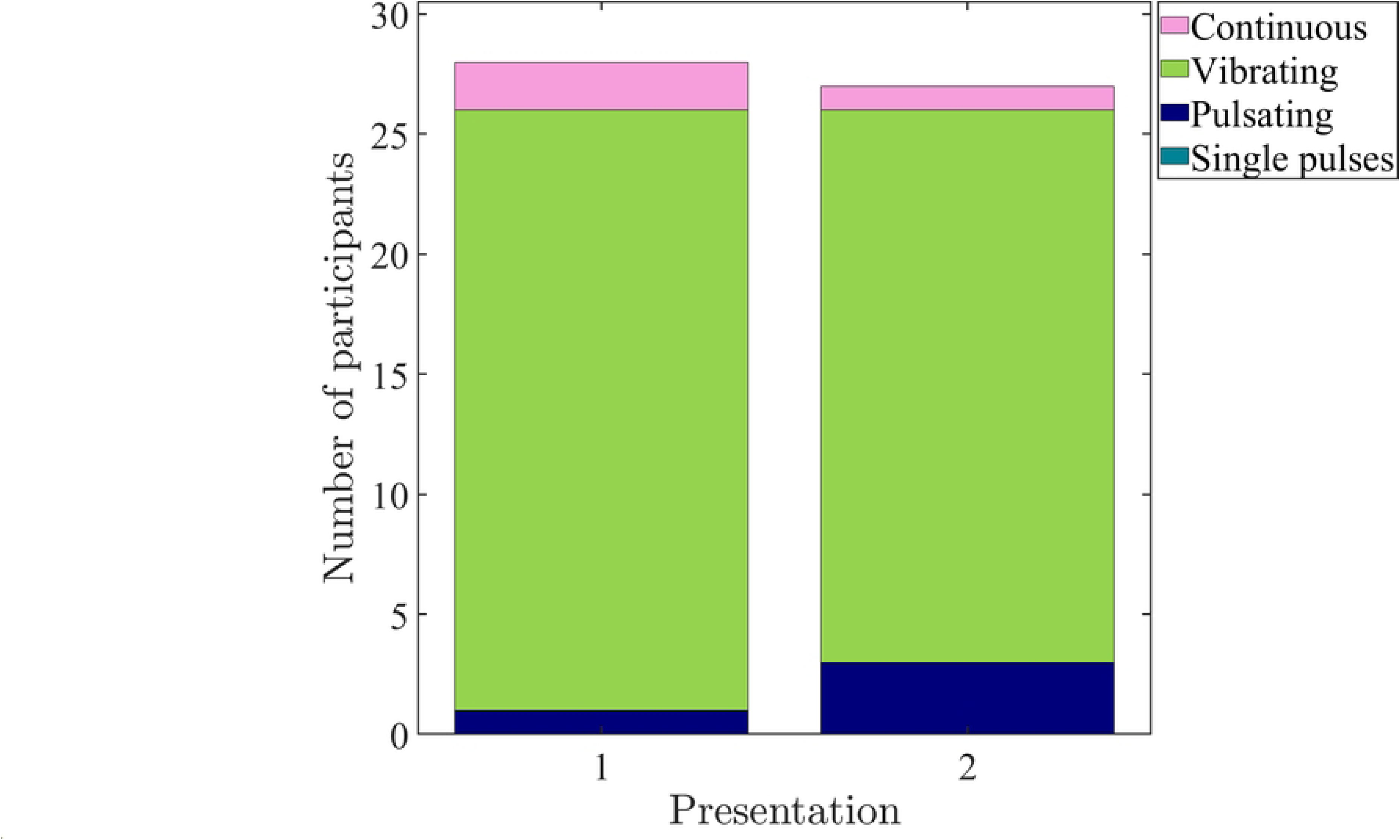
Mean relative proportion of muscle twitch location of study 2 during the work tasks. The proportions are given at the warning and at the intolerance threshold, reported by the participant and documented by the independent observer, for the work tasks reading, operating a cordless screwdriver, and operating a polishing machine. The proportions are averaged over the participants who experienced muscle twitches.

## Discussion

In two studies, we evaluated a circumferential electrocutaneous warning signal during three work tasks: reading, operating a cordless screwdriver, and operating a polishing machine. In study 1 (*n* = 32) the pulse interval was determined individually for each participant, following our previous approach [6, 8–10], whereas in study 2 (*n* = 29) a fixed pulse interval of 36 ms was applied to all participants and the warning threshold was assessed in addition to the intolerance threshold. Electrocutaneous warning during work tasks proved feasible in both studies. The warning signal was perceived as a pulsating or vibrating sensation at rest by 89 % of the participants of study 2 without individual parametrization, the warning threshold of 8 mA did not differ between the three tasks, and the intolerance threshold was lower during reading (18 mA) than during screw-driving and polishing (20 mA each). In study 1 the intolerance threshold was likewise task dependent and highest during polishing, at overall lower amplitudes. In both studies the dominant location of muscle twitches was ventral, with a diversification of the location towards higher amplitudes.

The single-pulse reference thresholds of study 1 reproduce the hierarchy of perception, muscle twitch, and intolerance thresholds reported previously [5, 9, 34, 35], as well as the knocking sensation at the perception threshold [5, 9]. The difference between the single-pulse intolerance threshold, not reached within 25 mA for 27 of 32 participants, and the intolerance threshold of the warning pattern at medians of 14 mA to 16.5 mA is notable. The perceived magnitude of electrocutaneous stimulation increases with the number of pulses and with decreasing pulse interval at constant amplitude [36–39], which supports our decision to derive the amplitude from the pattern itself rather than from single-pulse thresholds.

The individual determination of the pulse interval used in our previous studies and in study 1 is precise but impractical for a wearable system. The re-evaluation of the individual pulse interval ranges of 80 participants [7] showed that no range for a vibrating sensation is shared by all participants, but that the combined ranges for a pulsating and a vibrating sensation overlap between 33 ms and 40 ms. Both temporal qualities produce comparable levels of alertness for a circumferential pattern [6]. With the resulting fixed value of 36 ms, 89 % of the participants of study 2 reported a pulsating or vibrating perception in both presentations at rest. Two participants in the first and one in the second presentation reported a continuous perception, which corresponds to a single sustained stimulus per electrode pair and may reduce the alerting character of the pattern [6, 7]. A fixed parametrization is therefore sufficient for the large majority of users, while an optional adjustment of the pulse interval should remain available. This applies to a binary warning. Where several stimulus levels have to be distinguished, as in prosthetic feedback that encodes continuous information, individual calibration to the just noticeable difference may be required [40].

The warning threshold of study 2 was 8 mA at rest and during all three tasks, so that the perception of the pattern is robust against both cognitive and motor load. The processing of task-irrelevant stimuli depends on the type of load, as high perceptual load reduces it whereas high load on cognitive control increases it [41]. That the warning threshold remained unchanged across conditions that differ in both respects suggests that the circumferential pattern is not processed like an arbitrary distractor. The intolerance threshold, in contrast, depended on the task in both studies. It was highest during polishing in study 1, and lowest during reading in study 2. The two tool-based tasks differ in the temporal structure of the mechanical load. The polishing machine runs continuously at 3000 revolutions per minute and is guided with both arms, whereas screw-driving produces short, intermittent loads separated by repositioning of the tool. We did not measure the vibration transmitted to the arm, so the following interpretation remains a hypothesis. Mechanical and electrical tactile stimuli are processed by partly overlapping afferent populations [42]. For mechanical stimuli, masking has been shown to depend on the receptor channel involved, as a masker raises the detection threshold of a test stimulus only when both fall within the frequency range of the same receptor system, whereas cross-channel masking does not occur [43]. Within a channel, the threshold shift increases with the sensation level of the masker [43, 44]. Whether a comparable interaction occurs between mechanical vibration and electrocutaneous stimulation cannot be decided from our data, as neither the vibration spectra of the tools nor the receptor populations addressed by the circumferential pattern were determined. Our previous finding of elevated electrosensory thresholds under a defined mechanical vibration [10] is consistent with such an interaction.

Masking alone, however, does not explain the dissociation between the two thresholds, which is also a difference to our previous results, where perception, attention, and intolerance thresholds all increased under a defined mechanical vibration [9, 10]. The attention threshold determined there is the closest analogue to the warning threshold, but it was obtained with a single pulse at one electrode pair, whereas temporal integration across pulses [36] and the sequential circumferential presentation [6] make the warning pattern considerably more salient. Masking acts primarily on the detection threshold, whereas suprathreshold intensity discrimination is affected mainly close to the masked threshold and becomes largely independent of the masker at higher sensation levels [45, 46]. The warning threshold, however, is a judgment of attention capture, and the intolerance threshold a judgment of tolerance, which task engagement plausibly shifts upwards. Separating these contributions requires a design in which detection, attention, and tolerance are assessed under the same task load.

The absolute intolerance thresholds were consistently higher in study 2 (18 mA, 20 mA, 20 mA) than in study 1 (14 mA, 14 mA, 16.5 mA). The most plausible explanation is the pulse interval and the charge applied per burst. At a pulse interval of 36 ms, 11 pulses are delivered per 0.4 s burst, compared with roughly 21 pulses at 19 ms, so that the charge applied per burst at a given amplitude is about half as large in study 2. As the pulse rate decreases, a longer pulse width would be required to maintain the same perceived intensity [38], and the perceived intensity increases with the applied charge and with the number of pulses [36, 39, 47]. Since the pulse width was fixed at 150 µs, a higher amplitude is required instead. The opposite sex composition of the two groups contributes as well, so that the comparison remains descriptive. In both studies a substantial proportion of participants tolerated the warning signal up to the maximum amplitude of the setup, so that the reported intolerance thresholds are right-censored and their variability is underestimated. The task dependence of the intolerance threshold is systematic but small compared with the differences between participants, so that the distributions in Fig 4 and Fig 12 overlap almost completely although the connecting lines show that the within-participant differences point consistently in one direction. The ceiling of the setup, 25 mA in study 1 and 24 mA in study 2, additionally suppresses the differences of the participants who reached it, so that the reported effect is likely to be underestimated rather than overstated. For the intended application this ceiling is less critical than it appears, because the relevant operating point is the warning threshold of 8 mA, which leaves a considerable usable operating window in all investigated tasks.

Muscle twitches occurred in both studies and remain a concern for a practical system. In study 1 the amplitude was ramped up to the intolerance threshold, so that the twitch reports refer to the whole amplitude range and are comparable to the intolerance threshold of study 2. On this basis the proportions were similar, with 88 %, 81 % and 84 % of the participants in study 1 and 86 %, 68 % and 71 % in study 2. For the intended application, however, the system is operated at the warning threshold, where only 7 % of the participants of study 2 reported muscle twitches. The strongly different sex composition of the two groups is a further plausible contribution, as women have been reported to show fewer muscle twitches during electrocutaneous stimulation [10] and differ in muscle fibre composition and contractile kinetics [48, 49].

The dominant location of the muscle twitches was ventral in both studies, and in study 2 it diversified towards dorsal, lateral, and whole-arm responses at the intolerance threshold, which we attribute to the spread of the electric field and the recruitment of additional motor units at higher amplitudes [50, 51]. The two studies differ, however, with respect to the medial region, which accounted for 28 % to 35 % of the reported twitches in study 1 but did not occur at all at the intolerance threshold during the work tasks in study 2, while it accounted for 4 % and 15 % of the participant reports at rest. Our previous parameter studies found muscle twitches most often at the medial electrode pairs [5, 6, 9], which is plausible given the superficial course of the median and ulnar nerves at the medial aspect of the upper arm [52], so that study 2 is the deviating case. Sex-related differences in the tissue between the electrodes and the excitable structures may contribute as well, and they act in opposite directions for the two relevant structures. The skin itself is thinner in women [53], so that the cutaneous afferents lie closer to the electrodes, which is consistent with the lower perception thresholds reported for women [10, 54]. The subcutaneous fat layer at the upper arm, in contrast, is thicker in women, so that the combined thickness of skin and subcutaneous tissue is greater [55] and the underlying motor structures are farther from the electrodes, which is consistent with the lower incidence of muscle twitches [10]. This effect would act most strongly medially, where the twitch-prone structures lie deeper than ventrally. A methodological cause is equally possible, as study 1 documented the location as electrode pair numbers with a subsequent mapping onto regions, whereas in study 2 the region was selected directly. In addition, the medial region is the least accessible to visual inspection, so that the observer reports of study 2 are likely to underrepresent it.

For a wearable warning system, involuntary arm movements are a potential hazard of their own, even though they increase the salience of the signal. During reading, 19 % of the participants of study 1 showed arm movements. In study 2, one participant showed a pronation of the lower right arm and two participants briefly let go of the polishing machine. Operating the system close to the warning threshold is the most direct mitigation. Beyond that, our comparison of TENS and textile electrodes showed less frequent muscle twitching for the smaller and more circular textile electrodes within a cuff [9], and the direction of current stimulation offers a further, although limited, degree of freedom [8], so that optimization of electrode geometry and placement remains the most promising route.

Participant self-report and independent observation were obtained independently, as the observer documented the twitches during the presentation without commenting on them and the participant was asked only afterwards. The two assessments agreed in the binary distinction between twitch and no twitch, but differed in the classification of the twitch strength in a task-specific manner. During reading, the observer documented more arm movements than the participants reported, whereas during the two tool-based tasks the relation was reversed. During reading, attention directed at the text plausibly reduces the awareness of involuntary movements of the otherwise resting arm, whereas during the tool-based tasks the superimposed tool movement impedes their visual detection by the observer. Both methods remain subjective, and an objective acquisition by surface electromyography [56] should be included in future studies.

Our studies have several limitations. Both study groups were young and homogeneous in age (29 *±* 7 years and 28 *±* 8 years), so that age-related differences in electrocutaneous perception could not be assessed [54, 57], and the sample sizes of 32 and 29 participants limit the statistical power [58]. The stepwise variation of the amplitude and the ceiling of the setup produce a large number of zero differences, which reduces the effective sample size of the paired comparisons. In study 2, only 13, 17 and 15 of the 29 participants contributed a non-zero difference to the three comparisons of the intolerance threshold. The sex distribution was unbalanced in opposite directions in the two studies, which limits their comparability. Both groups were recruited in a university setting, which restricts the generalizability to working professionals. Four participants took part in both studies, which may have introduced a learning effect for these individuals. The maximum amplitude of the setup was reached by a relevant proportion of participants, so that the intolerance thresholds are right-censored. As stated above, the charge density of the present setup remains far below the established limits, so that future studies might employ a higher maximum amplitude [29, 30].

The application context was only approximated. The warning threshold was first determined at rest, where the participants could direct their full attention to the stimulation, and the work tasks reduce this limitation but remain simplified. The vibration transmitted to the arm was not measured and is expected to be low compared with heavy equipment, for which reported hand-arm vibration values reach 8.9 m/s^2^ even for hand-guided tools [59]. As the tasks were presented in the same fixed order in both studies, sequence effects cannot be excluded.

The resolution of the two central measures was limited by the protocol. The warning threshold was determined in steps of 2 mA, so that differences below this step size between the tasks could not be resolved, which qualifies the finding that the warning threshold did not depend on the task. In study 1, the individual determination of the pulse interval started at 19 ms and the interval was only adjusted when the sensation was not described as vibrating, so that the reported median of 19 ms is influenced by this starting value.

The measurement itself has further limitations. Thresholds were determined by a stepwise increase of the amplitude and depend on the subjective report of the participants. For the warning threshold, an adaptive psychophysical procedure would have provided a more precise estimate [60], whereas for the intolerance threshold repeated approaches from both sides are not feasible, as the stimulation has to be stopped as soon as the threshold is reached. Neither study was blinded and the operator was aware of the hypotheses, so that an experimenter expectancy effect cannot be excluded [61]. The two studies also differ in their documentation of muscle twitches, as study 1 recorded the location as electrode pair numbers and used the strength category “perceptible”, which was omitted in study 2 because it cannot be verified by an external observer. In study 2, the observer could not inspect all regions of the upper right arm equally well, as the medial region faces the trunk and is difficult to see while the participant is seated or operating a tool, so that muscle twitches at this location may be underrepresented in the observer reports.

The setup is not yet a wearable system. The self-adhesive TENS electrodes used in both studies are not suitable for a wearing time of eight hours in everyday work, because the hydrogel layer degrades during use and requires regular replacement [62]. In addition, the laboratory setup is not portable, so that the warning cannot yet be evaluated in a real work environment.

## Conclusion

Our studies contribute to the development of an electrocutaneous warning system for workers in potentially hazardous situations. For the first time, we applied a circumferential electrocutaneous warning signal during representative work tasks, once with an individually adjusted pulse interval and once with a fixed pulse interval that was identical for all participants.

We demonstrate that electrocutaneous warning during work tasks is feasible and that a fixed parametrization is sufficient for the large majority of users when a binary warning is to be conveyed, as 89 % of the participants perceived the warning signal at rest in the intended pulsating or vibrating quality. The warning threshold was 8 mA and was independent of the task, whereas the intolerance threshold was lower during reading (18 mA) than during screw-driving and polishing (20 mA each). With individually adjusted pulse intervals, the intolerance threshold was likewise task dependent and highest during polishing, at overall lower amplitudes. Muscle twitches were largely absent at the warning threshold and became frequent towards the intolerance threshold, with a ventral dominance of the location in both studies and a diversification at higher amplitudes.

Several conclusions follow for the design of a wearable warning system. An individual determination of the pulse interval is not required for most users, but an optional adjustment should remain available for individuals reporting a continuous perception. Should the system later encode several distinguishable levels, individual calibration might become necessary again. Electrode optimization, in particular the textile electrode cuff investigated in our previous work [9], remains the most promising route to further reduce muscle twitches and to achieve a wearing time compatible with a full workday. Objective measurement of motor responses, for example by surface electromyography, should complement the observer-based assessment.

Future work will address longer exposure times in order to quantify adaptation and fatigue effects, work tasks with stronger vibration exposure, further disturbances such as noise, and individual factors such as age and skin properties in a larger and more diverse participant group covering the working age range. Field tests in industrial work environments with a portable setup are required to assess the effectiveness of the warning under practical conditions. Beyond occupational safety, the findings may also be relevant for other applications of electrocutaneous stimulation, such as sensory feedback in prosthetics [40, 63, 64].

## Supporting information

**S1 PDF. Story of the reading task: “The Disappeared Professor”.**

