## Supplemental Data 1 for "Individual versus fixed parametrization of an electrocutaneous warning signal during manual work tasks"

### The Disappeared Professor

It was a completely normal Monday morning at the university. The sky was grey, the coffee in the cafeteria as weak as always, and most students had barely slept. Still, they were all seated on time at 8 a.m. in Lecture Hall 3, waiting for Professor Weber.

He was known for his exciting way of teaching. He explained dry subjects like statistics as if they were crime stories. Nobody wanted to miss his lectures.

But today, something was different.

The lectern was empty. No laptop. No papers. No professor.

Only a white sheet of paper lay there. Written in black marker were the words:

*“The truth lies in Room 404. Follow the clues.”*

A murmur went through the room. “Does Room 404 even exist?” someone asked. “Isn’t that the internet error?” another laughed. But no one really knew whether the note was a joke—or a new teaching method.

Some wanted to leave. Others were curious. In the end, five students stood up. They wanted to find out what was behind it.

There was Lara, who always wanted to know everything. Jonas, who rarely spoke but noticed everything. Malik, who loved math and riddles even more. Sophia, who always carried a camera because she wanted to document everything. And Daniel, who really didn’t have time—but loved adventure.

They left the lecture hall and began their search.

### Clue 1 – The Library

The five went to the library. On the way, they wondered what "Room 404" could mean. “Maybe it’s a hidden room? Or a code?” asked Lara. Jonas said nothing, but he had pocketed the note.

At the library entrance, they saw a shelf with old books. Between two dusty encyclopedias was a small envelope. On it was written:

*“Knowledge is the key. Look where no one speaks aloud.”*

They entered the reading room. It was quiet there. Most seats were empty. In the corner stood a large wooden globe. Malik discovered a small metal box underneath it.

Inside the box was a USB stick. They plugged it into Sophia’s laptop. A video file opened: Professor Weber!

*“Congratulations. You’ve taken the first step. This is not a game – it’s a test. No grades, no formulas. Only teamwork and thinking. Find the next location: where the light never goes out.”*

### Clue 2 – The Server Room

“Where does the light never go out?” Lara thought aloud. Jonas pointed at the basement map. “Server room,” he said quietly.

They asked the janitor, who looked a little suspicious but eventually handed over the key. The server room was cool and full of quiet humming noises. Between two cabinets hung a small note:

*“Statistics is not dry. It is hidden. 3 – 14 – 27.”*

“Those are page numbers!” Malik exclaimed. “In his own book!”

They ran back to the library. On page 3 was a QR code. On page 14, a photo of a door. On page 27, there was only one word: **“Attic.”**

### The Finale – The Attic

They found the key at the reception desk. The attic was dark and smelled of dust. Between old furniture and boxes sat—Professor Weber!

He grinned. “Welcome. You did it. You thought together, searched, connected the dots—and didn’t give up. That’s worth more than any formula.”

He stood up and handed them an envelope. “A bonus point. For each of you. And one question: Do you want to join again next time?”

The five laughed. Of course they did.
